# A FASII inhibitor prevents staphylococcal evasion of daptomycin by inhibiting phospholipid decoy production

**DOI:** 10.1101/427989

**Authors:** Carmen J. E. Pee, Vera Pader, Elizabeth V. K. Ledger, Andrew M. Edwards

**Author notes:** These authors contributed equally (Carmen J. E. Pee, Vera Pader, Elizabeth V. K. Ledger).

## Abstract

Daptomycin is a treatment of last resort for serious infections caused by drug-resistant Gram-positive pathogens such as methicillin-resistant *Staphylococcus aureus*. We have shown recently that *S. aureus* can evade daptomycin by releasing phospholipid decoys that sequester and inactivate the antibiotic, leading to treatment failure. Since phospholipid release occurs via an active process we hypothesised that it could be inhibited, thereby increasing daptomycin efficacy. To identify opportunities for therapeutic interventions that block phospholipid release, we first determined how the host environment influenced the release of phospholipids and inactivation of daptomycin by *S. aureus*. The addition of certain host-associated fatty acids to the growth medium enhanced phospholipid release. However, in serum, the sequestration of fatty acids by albumin restricted their availability to *S. aureus* sufficiently to prevent their use in the generation of released phospholipids. This finding implied that in host tissues *S. aureus* is likely to be completely dependent upon endogenous phospholipid biosynthesis to generate lipids for release, providing a target for therapeutic intervention. To test this, we exposed *S. aureus* to AFN-1252, an inhibitor of the staphylococcal FASII fatty acid biosynthetic pathway, together with daptomycin. AFN-1252 efficiently blocked daptomycin-induced phospholipid decoy production, even in the case of isolates resistant to AFN-1252, which prevented the inactivation of daptomycin and resulted in sustained bacterial killing. In turn, daptomycin prevented the fatty acid-dependent emergence of AFN-1252-resistant isolates. In summary, AFN-1252 significantly enhances daptomycin activity against *S. aureus* by blocking the production of phospholipid decoys, whilst daptomycin blocks the emergence of resistance to AFN-1252.

## Introduction

Daptomycin is a lipopeptide antibiotic of last resort used to treat infections caused by drug-resistant Gram-positive pathogens such as methicillin-resistant *S. aureus* (MRSA) and vancomycin-resistant enterococci (VRE) [1,2]. The target of daptomycin is the bacterial membrane, where it causes mis-localisation of enzymes required for cell wall biosynthesis, loss of membrane potential and integrity, and rapid bacterial death [1,3,4].

Resistance to daptomycin can arise spontaneously via mutations in genes associated with phospholipid or peptidoglycan biosynthesis [1,5,6]. However, whilst resistance has been reported to arise during treatment, it is a rare occurrence and does not explain why daptomycin treatment fails in up to 30% of cases [7,8]. In a bid to identify additional mechanisms by which *S. aureus* can withstand daptomycin treatment, we discovered that upon exposure to the antibiotic, *S. aureus* releases phospholipids into the extracellular space [9]. These phospholipids act as decoys, sequestering daptomycin and preventing it from inserting into the bacterial membrane. This decoy-mediated antibiotic inactivation led to treatment failure in a murine model of invasive MRSA infection, suggesting that it could affect daptomycin efficacy in patients [9]. Furthermore, the production of phospholipid decoys also occurs in enterococci and streptococci, suggesting a broadly conserved mechanism for resisting membrane-acting antimicrobials [10].

The ability of released membrane phospholipids to inactivate daptomycin can be compromised in *S. aureus* by the quorum-sensing-triggered production of small cytolytic peptides known as the alpha phenol soluble modulins (PSMα) [9]. These peptides appear to compete with daptomycin for the phospholipid and thereby prevent inactivation of the antibiotic [9]. Whilst this may appear paradoxical, many invasive infections are caused by *S. aureus* strains defective for PSMα production due to defects in the Agr quorum-sensing system that triggers expression of the peptides [11-13]. Furthermore, serum apolipoproteins inhibits Agr and sequesters PSMs, which would be expected to allow wild-type bacteria to inactivate daptomycin [14-17].

The mechanism by which daptomycin triggers phospholipid release is currently undefined. However, we have shown that it is an active process that requires energy, as well as protein, cell wall and lipid biosynthesis [9,10]. The requirement for fatty acid biosynthesis for phospholipid release is important because it raises the prospect of targeting this process to enhance daptomycin efficacy. We have shown previously that inhibition of the FabF component of the FASII fatty acid synthetic pathway, using the antibiotic platensimycin, completely blocked phospholipid release [9,10]. Whilst platensimycin is unsuitable as a therapeutic drug due to poor pharmacological properties, the FabI inhibitor AFN-1252 shows more promising characteristics and a pro-drug variant is currently undergoing phase 2 clinical trials [18,19]. However, despite excellent *in vitro* activity, the therapeutic value of inhibitors of fatty acid synthesis as mono-therapeutic agents has attracted much debate [20,21]. Several bacteria, including *S. aureus*, can utilise fatty acids present in the host to generate phospholipids [21-24]. Although wild-type *S. aureus* strains cannot fully substitute exogenous fatty acids for endogenous fatty acids synthesised via FASII, there is evidence that some clinical isolates have acquired mutations that enable them to fully bypass endogenous fatty acid biosynthesis by utilising host-derived fatty acids [22,25,26]. Furthermore, *in vitro* experimentation suggests that the acquisition of such mutations is dependent upon the presence of host-associated fatty acids, which means that the frequency at which resistance to AFN-1252 emerges *in vivo* may have been under-estimated [25,26]. As such, the long-term viability of fatty acid synthesis inhibitors, such as AFN-1252, as mono-therapeutic antibacterial drugs is unclear and their ability to block daptomycin-induced phospholipid release in the presence of exogenous fatty acids undetermined [20,21].

Therefore, the aims of this work were to understand how the availability of fatty acids in the host influences the production of phospholipid decoys and determine whether AFN-1252 could be used in combination with daptomycin to provide a viable approach to combatting MRSA infection.

## Results

### Exogenous fatty acids modulate daptomycin-induced phospholipid release

The release of membrane phospholipids in response to daptomycin occurs via an active process that requires *de novo* phospholipid biosynthesis [9]. Whilst *S. aureus* has an endogenous fatty acid biosynthetic pathway (FASII), it can also incorporate fatty acids from the host into membrane phospholipid production [21-24]. Therefore, it was hypothesised that host-derived fatty acids would contribute to the production of lipids required for daptomycin-induced phospholipid release.

To enable accurate measurements of phospholipid release, these experiments were done in TSB containing, or not, one of several different fatty acids found in normal human serum [27]. Although the concentrations of each lipid species in serum varies, we employed a single concentration of 20 μM to enable direct comparison. Furthermore, the *S. aureus ∆agrA* mutant was used to avoid the Agr system compromising daptomycin inactivation [9].

As reported previously, exposure of the *S. aureus* USA300 *∆agrA* mutant to daptomycin in the absence of exogenous fatty acids resulted in the release of phospholipids into the extracellular space (Fig. 1A) [9]. Supplementation of the TSB growth medium with linoleic acid had no effect on the rate or quantity of phospholipid released, whilst the presence of myristic or palmitic acids resulted in a small increase in the quantity of phospholipids released at the latest time point (Fig. 1A). By contrast, the presence of oleic or lauric acids significantly enhanced both the rate and quantity of phospholipids released, relative to TSB without fatty acids (Fig. 1A).

**Figure 1.**
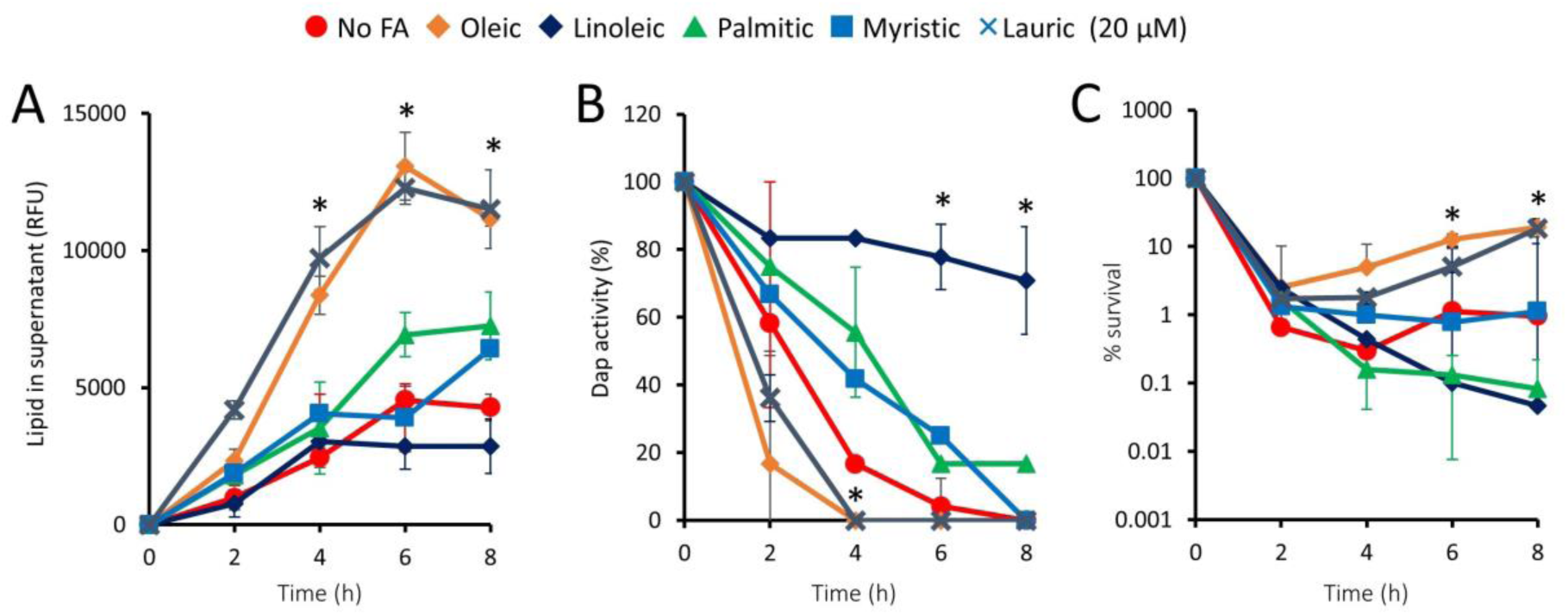
Effect of exogenous fatty acids on daptomycin-induced phospholipid release, daptomycin inactivation and bacterial survival. *S. aureus ∆agrA* was exposed to daptomycin (20 μg ml^−1^) in the presence of calcium (0.5 mM) and the indicated fatty acid supplements (20 μM) or none (No FA), and the release of phospholipids (A), antibiotic activity (B) and bacterial survival (C) measured over time. Data represent the means of 4 independent experiments and error bars show the standard deviation of the mean. Values significantly different (P <0.05) from bacteria in broth without fatty acid supplements were identified by 2-way repeated measures ANOVA and Dunnett’s post-hoc test (*).

As reported previously, in the absence of exogenous fatty acids, phospholipids released from the *∆agrA* mutant resulted in the inactivation of daptomycin (Fig. 1B) [9]. The increased release of phospholipids from bacteria incubated with oleic or lauric acids resulted in a slightly faster rate of daptomycin inactivation, whilst the presence of linoleic, palmitic or myristic acids reduced the rate of daptomycin inactivation (Fig. 1B). Of note, *S. aureus* failed to fully inactivate daptomycin in the presence of palmitic or linoleic acids, indicating that exogenous fatty acids can retard as well as promote the rate of phospholipid-mediated daptomycin inactivation (Fig. 1B).

In keeping with the effect of individual fatty acids on daptomycin inactivation, the presence of oleic or lauric acid promoted bacterial survival 10-fold above that seen for *S. aureus* incubated without fatty acids by 8 h. By contrast, the presence of palmitic or linoleic acids reduced survival approximately 10-fold, whilst myristic acid had no effect (Fig. 1C).

Taken together, these experiments demonstrated that certain fatty acids, such as oleic and lauric acids, can significantly enhance phospholipid release, whilst others are inhibitory or have no effect.

### Serum albumin restricts the utilisation of oleic acid by *S. aureus* for phospholipid release

Having established that fatty acids can modulate phospholipid release in TSB, we wanted to determine whether their presence in the host context had a similar effect. To do this, we firstly supplemented TSB with 50% delipidated human serum, which is deficient for fatty acids. Similarly to what was seen in TSB alone, exposure of the *∆agrA* mutant to daptomycin in TSB containing 50% delipidated human serum resulted in an initial fall in CFU counts, followed by a period of recovery (Fig. 2A). However, in contrast to our observations for TSB (Fig. 1C), the addition of oleic acid to TSB containing 50% delipidated serum had no effect on bacterial survival (Fig. 2A). In keeping with these data, the presence of oleic acid had no effect on the rate at which the bacteria inactivated daptomycin (Fig. 2B). This indicated that the ability of *S. aureus* to use oleic acid to promote phospholipid release was restricted by a factor found in serum but not TSB, although this was not quantified directly as serum proteins interfered with the dye-based assay system.

**Figure 2.**
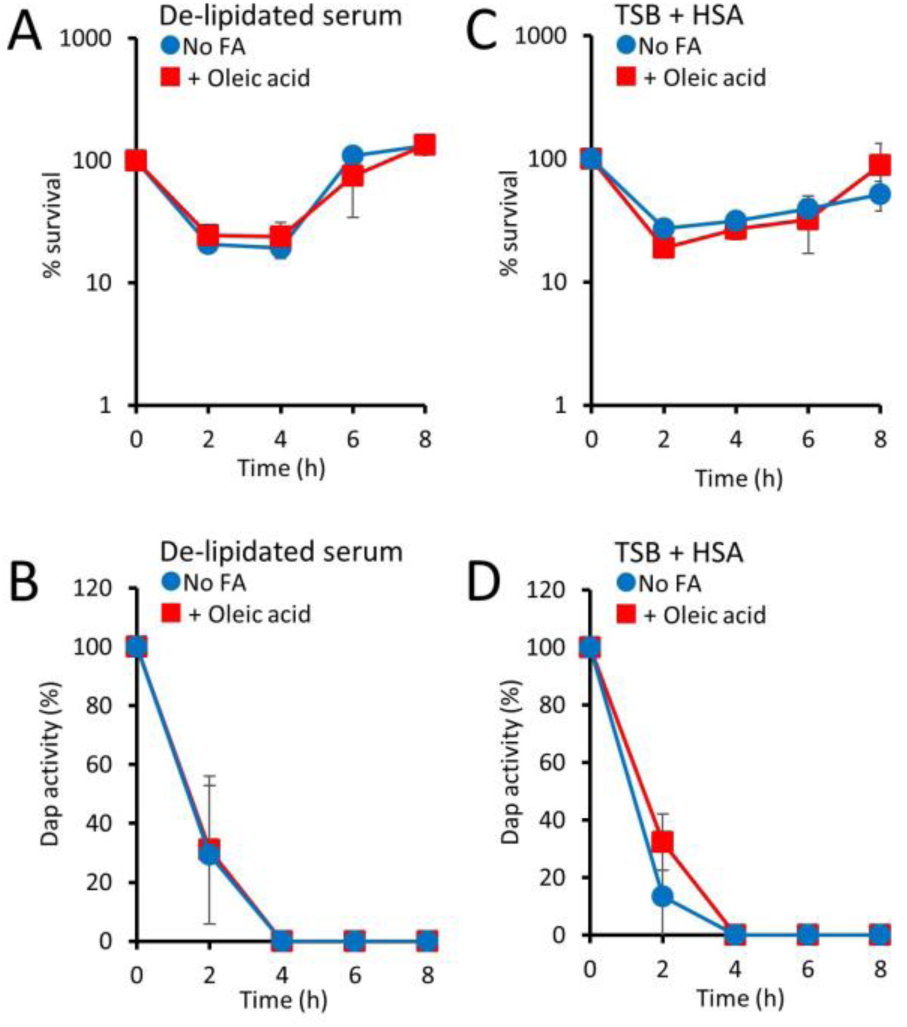
Human serum albumin prevents the use of exogenous oleic acid in daptomycin-induced phospholipid release. The *S. aureus ∆agrA* mutant was exposed to daptomycin (20 μg ml^−1^) in TSB containing 50% delipidated human serum and CaCl2 (0.5 mM) and supplemented with oleic acid (20 μM) or not (No FA), and bacterial survival (A) and antibiotic activity (B) measured over time. In a similar experiment, *S. aureus ∆agrA* was exposed to daptomycin in TSB containing human serum albumin (HSA) and CaCl2 and supplemented with oleic acid (20 μM) or not (No FA), and bacterial survival (C) and antibiotic activity (D) measured over time. Data represent the means of 4 independent experiments and error bars show the standard deviation of the mean. There were no significant differences in values obtained with oleic acid compared to un-supplemented medium (P >0.05) as determined by 2-way repeated measures ANOVA.

Fatty acids present in the bloodstream are typically bound to serum albumin, which acts as a carrier protein [28]. To determine whether the presence of this host protein restricted the availability of oleic acid for use in phospholipid release-mediated inactivation of daptomycin, the *S. aureus ∆agrA* mutant was exposed to daptomycin in TSB containing oleic acid and human serum albumin (HSA). By contrast to TSB only, the presence of HSA completely abrogated the increased rate of daptomycin inactivation and bacterial survival observed on supplementation with oleic acid, presumably due to sequestration of the fatty acid by the protein (Fig. 2C,D). Therefore, the sequestration of oleic acid by serum albumin prevents its use by *S. aureus* to promote daptomycin-induced phospholipid release.

### AFN-1252 blocks daptomycin-induced phospholipid release

The finding that HSA prevented the use of exogenous oleic acid by *S. aureus* to promote the rate of daptomycin inactivation indicated that this process is likely to be entirely dependent upon the FASII pathway *in vivo*. AFN-1252 is a FASII pathway inhibitor which blocks FabI and has shown potent activity against *S. aureus* in both pre-clinical and clinical testing [18,19]. Based on our previous findings [9], and the data described above, we hypothesised that AFN-1252 would enhance daptomycin activity against *S. aureus* by blocking the production of phospholipid decoys.

To test this, the *S. aureus ∆agrA* mutant was exposed to AFN-1252 (0.15 μg ml^−1^) in the absence or presence of daptomycin. Alone, AFN-1252 showed bacteriostatic activity (<10-fold drop in CFU counts after 8h) (Fig. 3A). As described previously, CFU counts of the *S. aureus ∆agrA* mutant exposed to daptomycin fell initially, before recovering due to the release of phospholipids that led to the inactivation of the antibiotic (Fig. 3A,B,C) [9]. However, when the *S. aureus ∆agrA* mutant was exposed to daptomycin in the presence of AFN-1252, there was a >500-fold drop in CFU counts, with no recovery of the bacterial population (Fig. 3A). Further analysis revealed that AFN-1252 almost completely blocked daptomycin-induced phospholipid release and the associated daptomycin inactivation (Fig. 3B,C), providing an explanation for the synergy observed when these antibiotics were used in combination.

**Figure 3.**
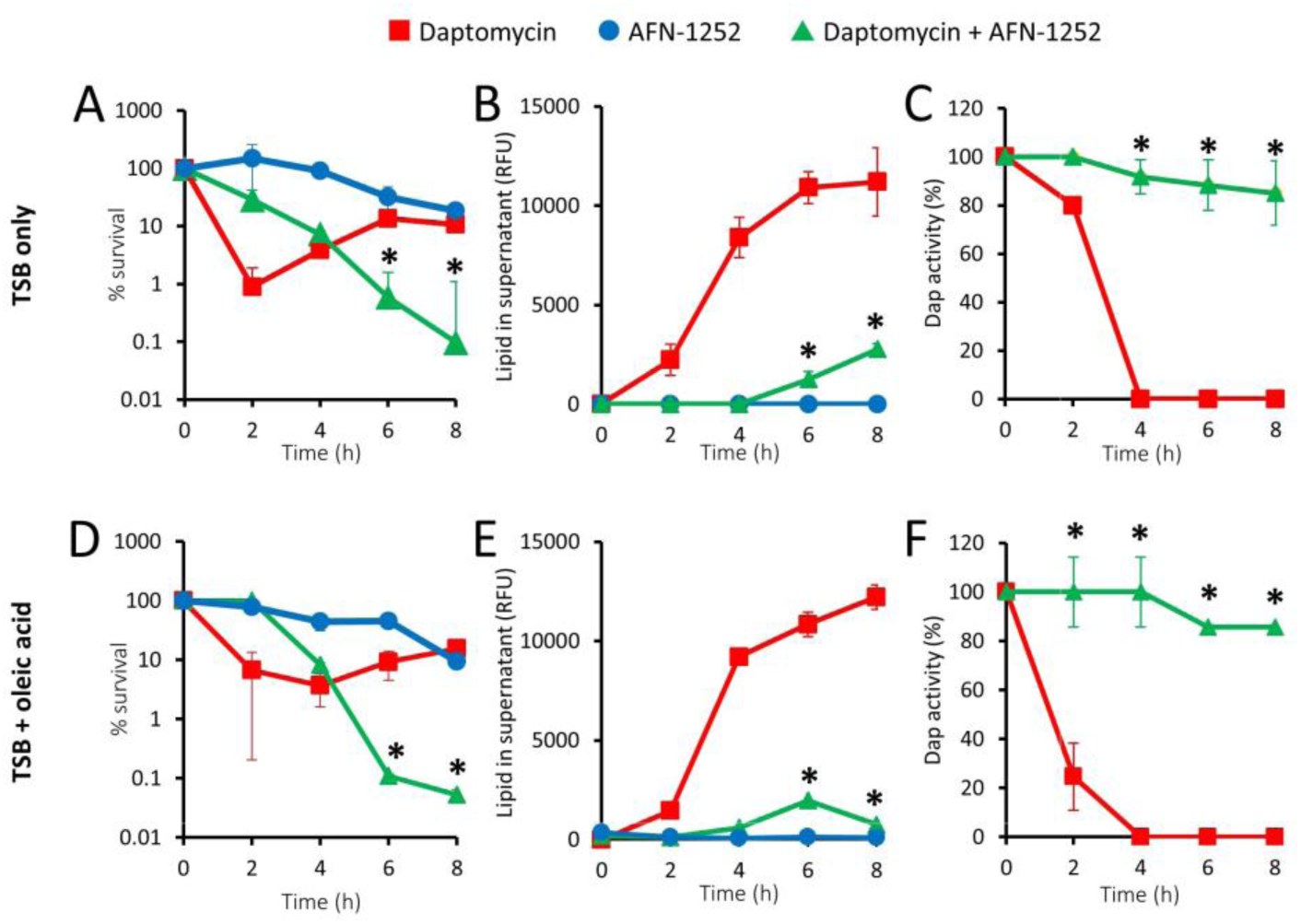
AFN-1252 blocks phospholipid release and therefore preserves daptomycin activity. The *S. aureus ∆agrA* mutant was incubated in TSB containing CaCl2 (0.5 mM) and daptomycin (20 μg ml^−1^) or AFN-1252 (0.15 μg ml^−1^), or both antibiotics in the absence (A,B,C) or presence (D,E,F) of oleic acid (20 μM). During incubation, bacterial survival (A,D), quantity of phospholipid released into the supernatant (B,E) and antibiotic activity (C,F) was measured over 8 h. Data represent the means of 4 independent experiments and error bars show the standard deviation of the mean. Values significantly different (P <0.05) from those obtained with bacteria exposed to daptomycin only were identified by 2-way repeated measures ANOVA and Dunnett’s post-hoc test (*).

Whilst our data indicated that HSA restricts the utilisation of fatty acids for phospholipid release (Fig. 2C,D), we considered the possibility that some unbound lipids may arise during infection because of damage to host tissues. Therefore, we repeated the experiments described in Figures 3A,B,C in the presence of oleic acid without HSA, which had previously been shown to significantly promote phospholipid release (Fig. 1A). The data generated from these experiments were almost identical to those from experiments done in the absence of the fatty acid (Fig. 3D,E,F). AFN-1252 showed clear synergistic activity when used in combination with daptomycin by blocking phospholipid release, even in the presence of unbound oleic acid (Fig. 3E). This resulted in the maintenance of daptomycin activity and a sustained killing effect on *S. aureus* (Fig. 3D,F).

Together, these data demonstrate that AFN-1252 prevents the production of phospholipid decoys, even in the presence of exogenous fatty acids which would otherwise enhance phospholipid release. Therefore, AFN-1252 prevents subsequent recovery of the population, enhancing the ability of daptomycin to kill *S. aureus*

### Exogenous fatty acids enable emergence of resistance to AFN-1252

The data described above indicated that the FASII inhibitor AFN-1252 in combination with daptomycin may be a promising therapeutic approach. However, it has been reported that *S. aureus* can acquire resistance to FASII inhibitors in the presence of exogenous fatty acids [25,26]. To confirm that these findings applied to AFN-1252, 10 parallel cultures of the USA300 *∆agrA* mutant were repeatedly challenged with AFN-1252 (0.15 μg ml^−1^) in the absence or presence of a physiologically relevant fatty acid cocktail as described previously [26]. Given the impact of HSA on daptomycin inactivation, parallel assays were done with or without the serum protein. After each exposure, bacterial susceptibility to AFN-1252 was determined by broth microdilution assays to establish the MIC.

As expected from previous reports, there was very little change in bacterial growth (Fig. 4A) or MIC (Fig. 4B) when *S. aureus* was repeatedly exposed to AFN-1252 in the absence of fatty acids [26]. However, in keeping with previous work, by the third round of exposure to AFN-1252 in the presence of fatty acids, with or without HSA, *S. aureus* was able to replicate in the presence of the antibiotic (Fig. 4A) [26]. The ability of *S. aureus* to grow in the presence of AFN-1252 after repeated exposure to the antibiotic in the presence of fatty acids, regardless of the presence of HSA, correlated well with data from subsequent MIC assays (Fig. 4C,D). When fatty acids were included in the MIC assays, there was a significant and large increase in the MICs of most cultures from 0.03125 μg ml^−1^ to more than 16 μg ml^−1^ (>512-fold) for bacteria that were exposed to AFN-1252 in the presence of exogenous fatty acids, regardless of the presence of HSA (Fig. 4C,D). Together, these data confirmed previous work showing that repeated exposure of *S. aureus* to AFN-1252 in the presence of exogenous fatty acids facilitated the emergence of fatty acid-dependent resistance to this antibiotic [26].

**Figure 4.**
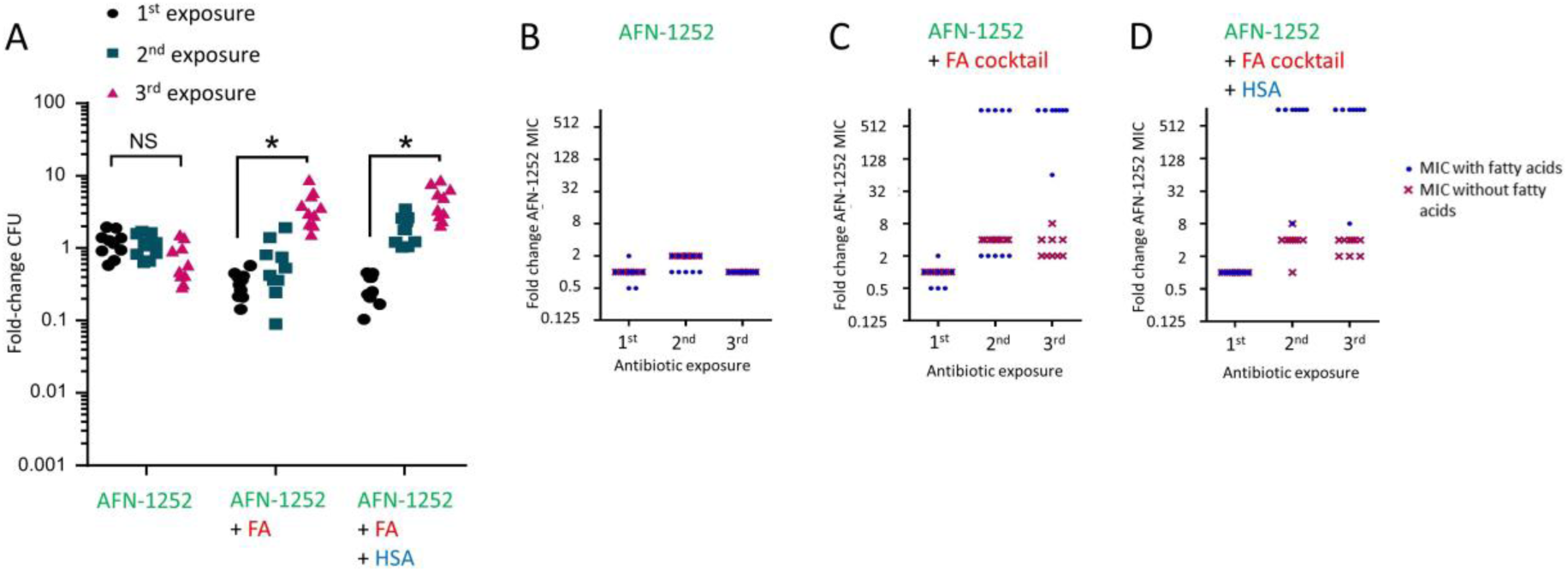
Exogenous fatty acids enable the acquisition of resistance to AFN-1252. Ten parallel cultures of the *S. aureus ∆agrA* mutant were exposed to AFN-1252 (0.15 μg ml^−1^) in the absence or presence of 50 μM fatty acids (FA) cocktail and absence or presence of HSA for 8 h before bacterial replication (A) and the AFN-1252 MIC determined in the absence or presence of FA cocktail (FA) (B,C,D). Each symbol represents an independent culture (n = 10 in each case). After 8 h exposure to AFN-1252, bacteria were recovered by centrifugation, washed and grown in antibiotic-free medium for 16 h before second and third rounds of antibiotic exposure and subsequent determination of bacterial survival and MIC. Differences in survival between the 1^st^ and 3^rd^ rounds of AFN-1252 exposure under identical conditions were analysed using a one-way ANOVA with Dunn’s multiple comparisons test (*P < 0.001).

### Daptomycin prevents fatty acid-dependent emergence of resistance to AFN-1252

Having confirmed that AFN-1252 resistance can arise in the presence of fatty acids, the next objective was to test whether combination therapy with daptomycin could prevent this. Therefore, the repeated antibiotic exposure experiment described above was re-run in the presence of daptomycin (20 μg ml^−1^) and in the absence or presence of exogenous fatty acids and HSA. As expected from previous data (Fig. 3), bacterial killing with daptomycin/AFN-1252 combination therapy was highly effective for the first two exposures, where bacterial survival was 1% or less after 8 hours. An increase in bacterial survival was observed on the third exposure, but bacterial growth was still inhibited with CFU counts not exceeding that of the original inoculum (Fig. 5A). Furthermore, this increase in survival was independent of the presence of fatty acids (Fig. 5A).

**Figure 5.**
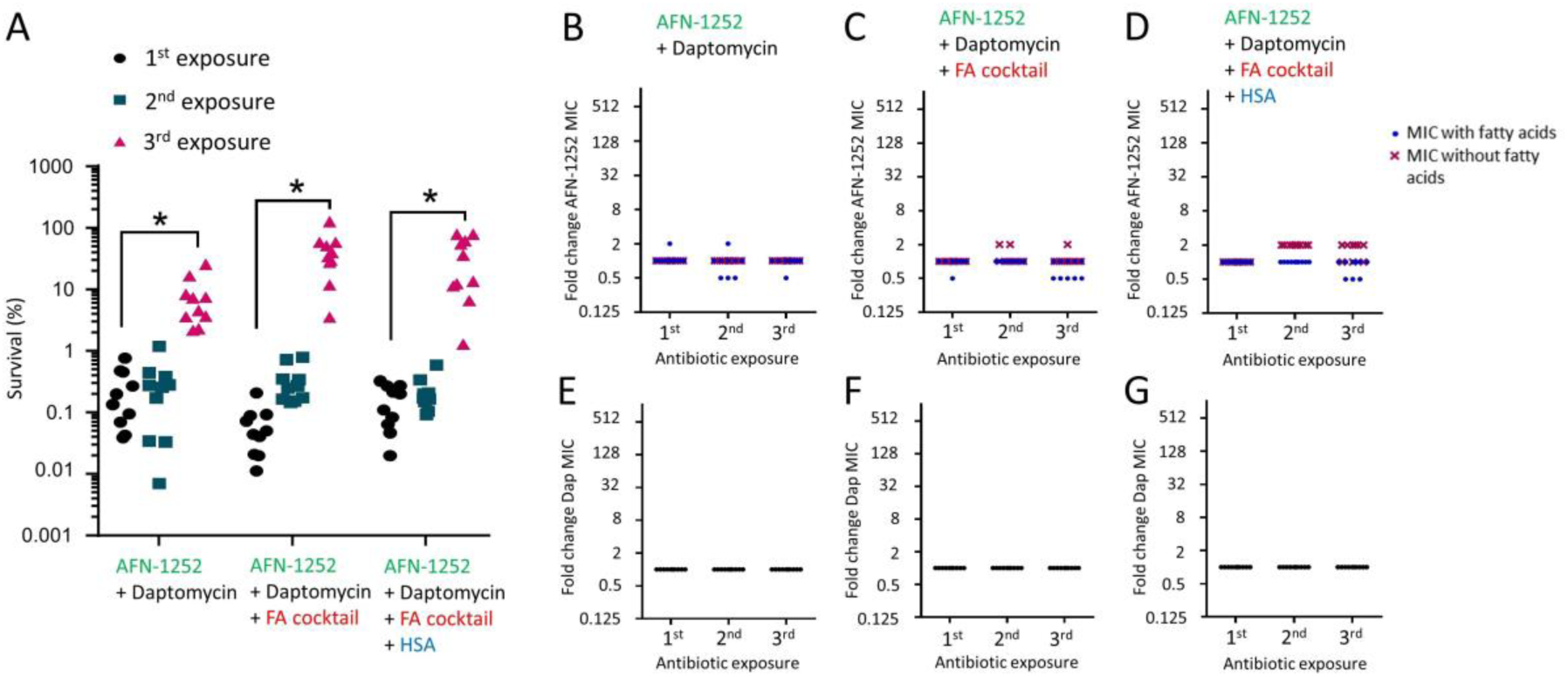
Daptomycin prevents the acquisition of fatty-acid enabled resistance to AFN-1252. Ten parallel cultures of *S. aureus ∆agrA* were exposed to daptomycin (20 μg ml^−1^) AFN-1252 (0.15 μg ml^−1^) in the absence or presence of fatty acid cocktail and absence or presence of HSA for 8 h before bacterial survival (A) and the AFN-1252 MICs determined in the absence or presence of fatty acids (B,C,D). The daptomycin MICs were also determined (in the absence of fatty acids) (E,F,G). After 8 h exposure to daptomycin and AFN-1252, bacteria were recovered by centrifugation, washed and grown in antibiotic-free medium for 16 h before second and third rounds of antibiotic exposure and subsequent determination of bacterial survival and MIC. Each symbol represents an independent culture (n = 10 in each case). Differences in survival between rounds of antibiotic exposure under identical conditions were identified using a one-way ANOVA with Dunn’s multiple comparisons test (*P < 0.001).

To determine whether daptomycin prevented the emergence of resistance to AFN-1252, MICs were determined by broth microdilution. By contrast to experiments with AFN-1252 alone, repeated exposure of *S. aureus* to AFN-1252 in the presence of daptomycin did not lead to an increase in MIC of the FASII inhibitor, even in the presence of fatty acids (Fig. 5B,C,D). Neither was there any increase in the daptomycin MIC (Fig. 5E,F,G). Together, these data demonstrate that daptomycin prevented the emergence of fatty acid-dependent resistance to AFN-1252 when the two antibiotics were used in combination.

Despite the increase in bacterial survival on the third exposure, this did not exceed the original inoculum (Fig. 5A), and the unchanged MIC values (Fig. 5B,C,D,E,F,G) indicated that AFN-1252 and daptomycin still had bacteriostatic activity. It is therefore likely that this increase in survival after 3 exposures was due to the acquisition of tolerance to daptomycin, which is consistent with a previous study [29].

### AFN-1252 blocks daptomycin-induced phospholipid release in AFN-1252-resistant strains

Having established that the combination of daptomycin and AFN-1252 prevented the emergence of AFN-1252 resistance, we next wanted to understand the underlying mechanism. To determine this, individual colonies of AFN-1252 resistant bacteria that had arisen in the presence of fatty acids and the presence or absence of HSA were picked. These were then assessed for survival, phospholipid release and ability to inactivate daptomycin in the absence or presence of fatty acids by comparison with the *∆agrA* mutant that had not been exposed to antibiotics.

As described above (Fig. 3), two independent colony picks of the *∆agrA* mutant that had not previously been exposed to antibiotics survived exposure to daptomycin by releasing phospholipids that completely inactivated the antibiotic (Fig. 6A,B,C). However, the presence of AFN-1252 increased the bactericidal activity of daptomycin by preventing phospholipid release and thus preserving the activity of the lipopeptide antibiotic, regardless of the presence of fatty acids (Fig. 6A,B,C).

**Figure 6.**
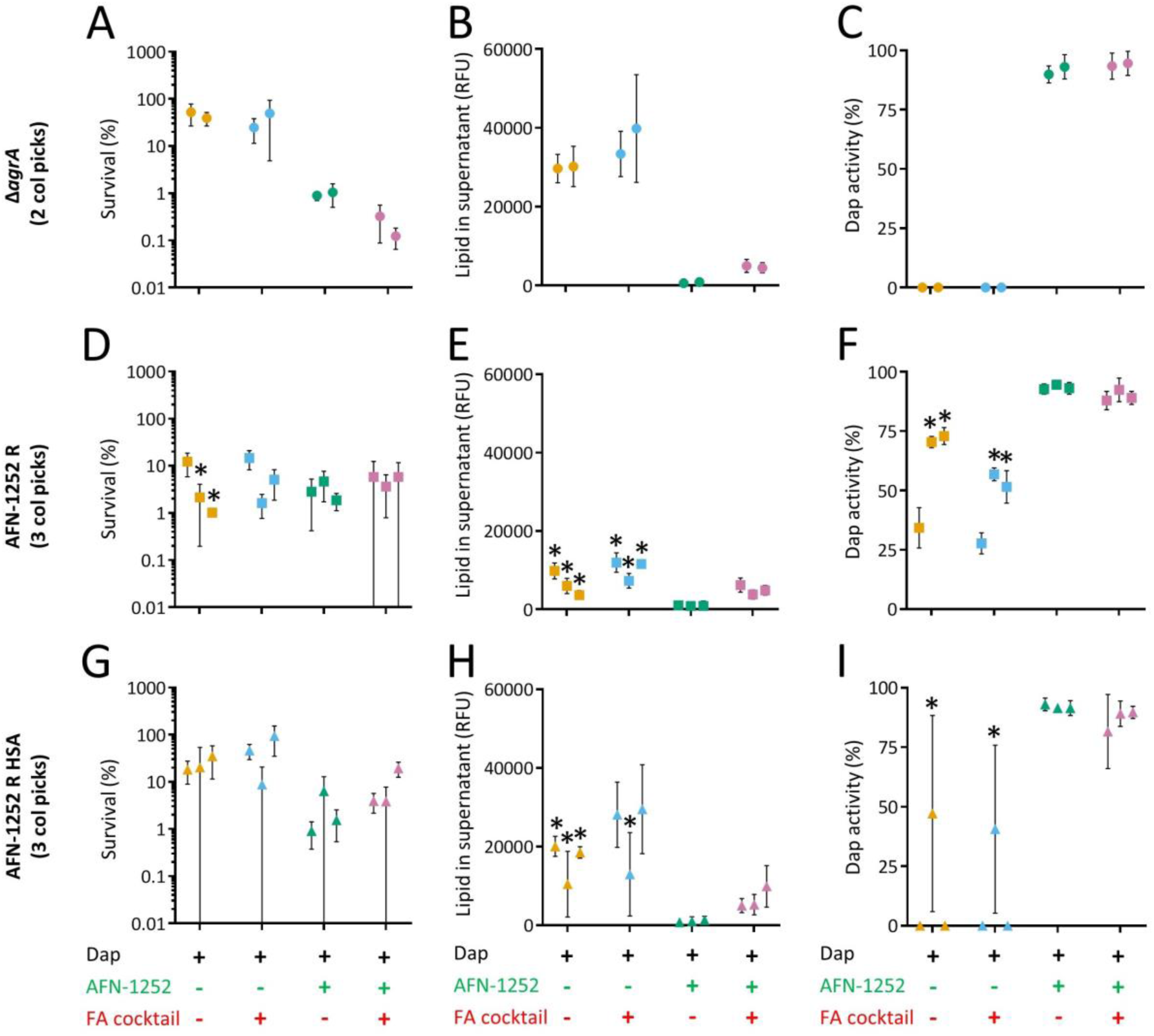
AFN-1252 prevents daptomycin-induced phospholipid release, even in the case of AFN-1252 resistant strains. Two independent isolates (each represented by a single circle) of the *S. aureus ∆agrA* mutant (USA300 *∆agrA)* that had not been exposed to antibiotic (A,B,C), three independent isolates of *S. aureus ∆agrA* that had acquired resistance to AFN-1252 in the presence of the FA cocktail but absence of HSA (AFN-1252 R) (D,E,F) or three independent isolates of *S. aureus ∆agrA* that had acquired resistance to AFN-1252 in the presence of the FA cocktail and HSA (AFN-1252 R HSA) (G,H,I) were exposed to daptomycin (Dap) in the presence or absence of various combinations of AFN-1252 (AFN) and fatty acid cocktail (FA) for 8 h. After this time, bacterial survival (A,D,G), the quantity of released phospholipid (B,E,H) and the activity of daptomycin (C,F,I) was determined. Data represent the mean of 3 independent experiments and error bars represent the standard deviation of the mean. Differences in survival, phospholipid release or daptomycin activity were compared between the AFN-1252 sensitive USA300 *∆agrA* isolates and AFN-1252 resistant isolates using a one-way ANOVA with Dunn’s multiple comparisons test (*P < 0.01).

Next, we assessed the survival of bacteria from 3 independent cultures that had acquired resistance to AFN-1252 during exposure to the antibiotic in the presence of fatty acids but not HSA (AFN-1252 R). Of these 3 isolates, 2 were more susceptible to daptomycin than the *∆agrA* mutant, apparently because they released lower levels of phospholipids that failed to fully inactivate the lipopeptide antibiotic (Fig. 6D,E,F). The remaining isolate reduced daptomycin activity by 70%, explaining its enhanced survival in the presence of daptomycin relative to the other 2 isolates. However, the presence of AFN-1252 completely abolished the ability of any of these isolates to inactivate daptomycin, even when exogenous FAs were present (Fig. 6D,E,F).

We then examined *S. aureus* isolates from 3 independent cultures that had acquired resistance to AFN-1252 during exposure to the antibiotic in the presence of fatty acids and HSA (AFN-1252 R HSA). Survival of these three AFN-1252-resistant isolates after exposure to daptomycin alone was not significantly lower than that seen for the AFN-1252-sensitive *∆sagrA* mutant. This was due to the release of sufficient phospholipid to inactivate all or most of the daptomycin that the bacteria were incubated with (Fig. 6G, H, I). However, despite the ability of these bacteria to grow in the presence of AFN-1252 when exogenous FAs were available, the FASII inhibitor almost completely blocked daptomycin-induced phospholipid release from all three isolates, even when the FA cocktail was present (Fig. 6G,H,I).

Together, these data reveal that fatty acid-enabled AFN-1252 resistance results in a reduced ability to release phospholipids in response to daptomycin (Fig. 6E, H). Furthermore, although these strains were deemed resistant to AFN-1252, daptomycin-induced phospholipid release was inhibited by the FASII inhibitor, even in the presence of exogenous fatty acids (Fig. 6F,I). This provides additional evidence that daptomycin-induced phospholipid release is dependent upon endogenous, FASII-mediated fatty acid biosynthesis, even in the case of AFN-1252 resistant bacteria that have access to exogenous fatty acids. As such, daptomycin-induced phospholipid release is efficiently blocked by AFN-1252, preventing inactivation of the lipopeptide antibiotic.

## Discussion

The high rate of daptomycin treatment failure warrants efforts to identify new approaches to enhance therapeutic outcomes. In this report we provide evidence that combining daptomycin with the fatty acid biosynthesis inhibitor AFN-1252 provides synergistic activity against *S. aureus* and reduces the frequency of drug resistance.

It is increasingly clear that the host environment modulates the susceptibility of bacterial pathogens to antibiotics due to the scarcity of nutrients and the induction of stress responses that result in changes in bacterial physiology [30,31]. Serum contains high concentrations of fatty acids, which can be exploited by *S. aureus* to produce phospholipids, reducing the metabolic costs associated with membrane biogenesis [21,23]. In keeping with this, we found that the presence of specific exogenous fatty acids, such as oleic or lauric acids, enhanced phospholipid release in response to daptomycin. However, *S. aureus* has strict requirements for the type of fatty acids that it can incorporate and, at least for wild-type strains, each phospholipid must have at least one fatty acid tail synthesised endogenously via FASII [32]. This requirement for FASII-mediated fatty acid biosynthesis to generate phospholipids was underlined by the ability of AFN-1252 to completely block phospholipid decoy release, regardless of the presence of oleic acid [32]. This provides evidence that daptomycin/AFN-1252 combination therapy would not be compromised by the availability of fatty acids in the host.

Whilst some exogenous fatty acids can be used for phospholipid biosynthesis during staphylococcal growth, it appears that their contribution to daptomycin-induced phospholipid release is severely compromised by the presence of serum albumin, which sequesters the fatty acids [28]. As described above, there is clear evidence that *S. aureus* can partially substitute endogenous fatty acid biosynthesis for exogenous host-derived fatty acids in the generation of phospholipids. However, our data demonstrate that the presence of serum albumin reduces the efficiency of this process sufficiently to prevent their use in daptomycin-induced phospholipid release, which must occur rapidly if the bacteria are to survive.

The successful clinical development of AFN-1252 would be a welcome addition to the arsenal of anti-staphylococcal antibiotics. However, although wild-type bacteria are dependent upon the endogenous FASII pathway to generate fatty acids for phospholipid biosynthesis, there is evidence that this is not the case in strains that have acquired mutations within the *acc* or *fabD* lipid biosynthetic gene loci [25,26]. These mutants can bypass FASII-mediated fatty acid production, conferring resistance to AFN-1252 in the presence of exogenous fatty acids [25,26]. It has been suggested that FASII bypass could compromise the long-term therapeutic viability of FASII inhibitors such as AFN-1252, a view that is supported by the identification of clinical isolates that are able to resist AFN-1252 in the presence of exogenous fatty acids [25]. However, early clinical studies have shown that AFN-1252 can successfully treat skin and soft tissue infections, albeit in a relatively small number of patients [18]. Therefore, it remains to be seen whether resistance to AFN-1252 becomes a significant clinical problem. However, given the ability of *S. aureus* to acquire resistance to antibiotics, it seems prudent to develop therapeutic strategies to prevent or overcome the emergence of resistance to AFN-1252. Our data provide support for the concept of AFN-1252 resistance via fatty acid dependent FASII bypass, but also demonstrate that it can be prevented by the presence of daptomycin, at least *in vitro*.

The combination of AFN-1252 and daptomycin could be described as a mutually-beneficial pairing; whilst AFN-1252 promotes daptomycin activity by blocking phospholipid release, daptomycin enhances AFN-1252 efficacy by preventing the emergence of resistance. This finding contributes to our growing appreciation for the potential of combination therapy approaches to circumvent resistance mechanisms. A well-established example of this is the combination of daptomycin and β-lactams that target penicillin-binding protein (PBP) 1. In this combination, daptomycin sensitises MRSA to β-lactam antibiotics by reducing the quantity of PBP2a available, whilst β-lactams sensitise *S. aureus* to daptomycin by increasing binding of the lipopeptide antibiotic to the bacterial membrane [33-36]. This phenomenon, known as the see-saw effect, significantly promotes killing of *S. aureus* relative to each of the antibiotics individually and is currently being assessed as a therapeutic option in a clinical trial [37].

Although the combination of daptomycin and AFN-1252 prevented the acquisition of resistance to either antibiotic, we did observe the emergence of tolerance to the lipopeptide antibiotic after the third exposure to the drugs. The acquisition of daptomycin tolerance has been reported previously and was found to occur via increased expression of the *dltABCD* operon, although the mechanism by which this reduced susceptibility was unclear [29]. Crucially, however, whilst this tolerance phenotype reduces the ability of the daptomycin/AFN-1252 combination to kill *S. aureus*, the antibiotics are still able to inhibit bacterial growth.

In summary, the presence of AFN-1252 prevents the phospholipid-mediated inactivation of daptomycin by *S. aureus*, whilst daptomycin prevents the fatty-acid dependent emergence of resistance to AFN-1252. Therefore, we propose that the combination of AFN-1252 and daptomycin may have therapeutic value for the treatment of serious MRSA infections.

## Methods

### Bacterial strains and growth conditions

*Staphylococcus aureus* strains USA300 wild-type and *∆agrA* mutant [9] were grown in tryptic soy broth (TSB) or on tryptic soy agar (TSA). For some assays TSB was supplemented with fatty acids including oleic acid, linoleic acid, palmitic acid, myristic acid or lauric acid (all obtained from Sigma). Since the serum concentrations of these fatty acids vary from 2 μM (lauric acid) to 122 μM (oleic acid) [27], assays were done with a single concentration (20 μM) within this range. For some assays, HSA was included (10 mg ml^−1^) to sequester fatty acids [22]. Bacteria inoculated onto TSA plates were incubated statically at 37 °C for 15-17 hours in air unless otherwise stated. Liquid cultures were grown in 3 ml broth in 30 ml universal tubes by suspending a single colony from TSA plates, and incubated at 37 °C, with shaking at 180 RPM to facilitate aeration for 15-17 hours to stationary phase. Staphylococcal colony forming units (CFU) were enumerated by serial dilution in sterile PBS and plating of aliquots onto TSA. Bacterial stocks were stored in growth medium containing 20% glycerol at - 80 °C.

### Antibiotic killing kinetics

*S. aureus* was grown to stationary-phase in 3 ml TSB with shaking (180 RPM) at 37 °C in 30 ml universals as described above. Bacteria were subsequently adjusted to a concentration of ~ 1 × 10^8^ bacteria ml^−1^ in fresh TSB containing 0.5 mM CaCl_2_ before antibiotics were added at the following concentrations: daptomycin (20 μg ml^−1^, Tocris), AFN-1252 (0.15 μg ml^−1^, Medchemexpress). For some experiments, TSB was supplemented with 50% normal human serum (Sigma), human serum albumin or fatty acids as indicated. Cultures were then incubated at 37 °C with shaking (180 RPM) and bacterial viability determined by CFU counts from samples taken every 2 h for 8 h.

### Daptomycin activity determination

The activity of daptomycin during incubation with *S. aureus* was quantified as described previously [9,10]. A well of 10 mm was made in TSA plates containing 0.5 mM CaCl2, followed by the spreading of stationary phase wild-type USA300 LAC (60 μl, ~10^6^ ml^−1^ in TSB) across the surface. When AFN-1252 was used in assays, TSA was spread with *Streptococcus agalactiae* COH1 instead of *S. aureus* as this bacterium is naturally resistant to the FASII inhibitor but susceptible to daptomycin. Thereafter the plate was dried before the wells were filled with filter-sterilised culture supernatant. Plates were then incubated for 16 h at 37 °C before the zone of growth inhibition around the well was measured at 4 perpendicular points. To accurately quantify daptomycin activity, a standard plot was generated for the zone of growth inhibition around wells that were filled with TSB supplemented with range of daptomycin concentrations. This enabled the conversion of the size of the zone of inhibition into percentage daptomycin activity.

### Phospholipid detection and quantification

*S. aureus* membrane lipid was detected and quantified using FM-4-64 (Life Technologies) as described previously [9,10]. Bacterial culture supernatants (200 μl) were recovered by centrifugation (17,000 x *g*, 5 min) and then mixed with FM-4-64 to a final concentration of 5 μg ml^−1^ in the wells of clear flat-bottom microtitre plates with black walls appropriate for fluorescence readings (Greiner Bio-one). Fluorescence was measured using a Tecan microplate reader, with excitation at 565 nm and emission at 660 nm to generate values expressed as relative fluorescence units (RFU). Samples were measured in triplicate for each biological repeat. TSB with or without fatty acids was mixed with the FM-4-64 dye and used as a blank. The fluorescent readings were analysed by subtracting the values from the blank readings and plotted against time.

### Antibiotic resistance selection assay

Stationary phase *S. aureus* was inoculated at ~10^8^ CFU ml^−1^ into 3 ml TSB with 0.5 mM CaCl2 containing antibiotics as specified, for 8 h per exposure. Daptomycin (20 μg ml^−1^) and/or AFN-1252 (0.15 μg ml^−1^) were used singly or in combination. After 8 h, bacterial survival was determined by calculating the fold-change (for assays with the bacteriostatic AFN-1252 only) or percentage-change (for assays with the bactericidal antibiotic daptomycin) in CFU relative to the inoculum. For repeated antibiotic exposure, 1 ml was removed from each culture post-antibiotic exposure, centrifuged (3 min, 17,000 x *g)* and the resulting pellet washed once in TSB before resuspension in 100 μl TSB. This was used to inoculate 3 ml TSB before incubation for 16 h at 37 ^o^C with shaking (180 RPM) in the absence of antibiotics. Bacterial exposure to antibiotics was then repeated twice for a total of three repeated exposures. In some experiments, the broth was supplemented with a fatty acid cocktail prepared as follows: myristic, palmitic and oleic acid (all from Sigma-Aldrich) were made up to 100 mM in dimethyl sulfoxide (DMSO) as described previously [26]. Where used, the fatty acid cocktail was diluted 1 in 2000 in culture medium to obtain a final concentration of 50 μM. In some cases, TSB was also supplemented with human serum albumin (Sigma-Aldrich) at 10 μg ml^−1^.

### Determination of antibiotic minimal inhibitory concentrations

Antibiotic susceptibility was determined using the broth microdilution procedure as described previously [38] to generate minimal inhibitory concentrations for daptomycin and AFN-1252. Antibiotics were diluted serially in 2-fold steps in TSB containing 0.5 mM CaCl_2_ in a 96-well microtitre plate to obtain a range of concentrations. In some assays, a fatty acid cocktail (50 μM) was added to the broth as described above for the resistance selection assay. Stationary phase bacteria were added to the wells to give a final concentration of 5 × 10^5^ CFU ml^−1^ and the microtitre plates incubated statically in air at 37 °C for 18 h. The MIC was defined as the minimum concentration of antibiotic needed to inhibit visible growth of the bacteria [38]. For some assays, fold change in MIC was calculated relative to the MIC of the USA300 *∆agrA* mutant which had not been exposed to antibiotics.

## Acknowledgements

EVKL is supported by a Wellcome Trust PhD Studentship (203812/Z/16/Z). AME acknowledges funding from the Department of Medicine, Imperial College and from the Imperial NIHR Biomedical Research Centre, Imperial College London.

